# Electron Microscopy Analysis of hAPP-Induced Neurodegeneration in the Glomerular Network

**DOI:** 10.1101/444265

**Authors:** Anastasia E. Robbins, Katherine S. Lehmann, Li Bai, Leonardo Belluscio

**Affiliations:** Developmental Neural Plasticity Section, National Institute of Neurological Disorders and Stroke, National Institutes of Health, Bethesda, MD 20892, USA.

**Author notes:** Co-first authors (Anastasia E. Robbins, Katherine S. Lehmann).

## Abstract

Loss of smell is an early indicator of Alzheimer’s disease (AD), making the olfactory system an accessible model to study the effect of AD related proteins such as Amyloid Precursor Protein (APP). The regenerative capacity of the system further enables studies of circuit recovery after APP-induced degeneration. While the cellular effects of APP are well documented, little is known about its effects on brain circuits at the ultrastructural level. To study circuitry changes, we overexpressed humanized APP with familial AD mutations (hAPP) in olfactory sensory neurons and performed serial electron microscopy on olfactory bulb glomeruli from both control and hAPP expressing mice. We found that hAPP-expressing mice showed a striking decrease in glomerular connectivity along with widespread changes of subcellular structures. By then turning off hAPP expression for 6 weeks we tested the capacity of glomerular circuits to recover and found clear restoration of both connectivity and subcellular features, including an increase in post-synaptic density to above the control level. These data provide an important ultrastructural view of olfactory regions associated with AD and suggest that circuit recovery is possible in brain tissue that has experienced APP-induced neurodegeneration.

## Introduction

Alzheimer’s Disease (AD) is the 6th leading cause of death in America, and loss of olfaction is commonly one of the first symptoms of AD, making the olfactory system an intriguing model to study mechanism of disease (Bacon et al. 1998; Kim et al. 2018). The olfactory system is highly plastic due to lifelong olfactory sensory neuron (OSN) turnover. OSNs are part of a regenerating olfactory epithelium that sends axons directly to the olfactory bulb (OB) in the brain (Graziadei and Graziadei 1979). In the OB, OSNs form structures called glomeruli where they synapse with output neurons that project to higher brain regions as well as local interneurons (Graziadei, Levine, and Graziadei 1978). OSNs turnover and their associated plasticity provides an opportunity to study the capacity for network recovery from neurodegenerative diseases like AD. Mutations and subsequent misfolding of Amyloid Precursor Protein (APP) is a suspected cause of AD (Hardy and Selkoe 2002; Hunter and Brayne 2018). It has been shown in two mouse models of AD that the induction of mutated humanized APP (hAPP) (Jankowsky, Slunt, et al. 2005) in OSNs leads to rapid glomerular disruption, loss in axonal connectivity, and olfactory behavioral deficits due to hAPP-induced OSN cell death (Cheng et al. 2013). Interestingly, many of these structural and behavioral deficits are reversible providing both functional and partial anatomical recovery (Cheng et al. 2013) however, how underlying neural connectivity reflects this degeneration and recovery remains unknown.

Ultrastructure analysis of the olfactory system provided early insights into its connectivity as research primarily focused on establishing baseline cellular anatomy and describing the development of the olfactory epithelium and bulb. The ability of glomeruli to regenerate after OSN axotomy has also been described through EM studies on frogs (Graziadei and DeHan 1973), hamsters (Morrison and Costanzo 1989) and mice (Au, Treloar, and Greer 2002) although to a limited extent. These studies used discrete micrographs to study basic anatomical changes and did not examine overall connectivity or recovery. Furthermore, these studies used lesions introduced through mechanical or chemical disruption at distinct times to study regenerative potential, but little is known about the olfactory glomeruli in an ongoing degenerative state. A recent study used light microscopy-based reconstruction to show that glomeruli from Parkinson Disease patients exhibited disorganization and volume loss (Zapiec et al. 2017) but the cellular, molecular, and connectivity consequences of the degeneration could not be determined at light resolution, furthermore this type of human post-mortem tissue study cannot assess the ability of neural circuits to recover following insult, an important question to explore in the broader context of addressing degenerative disease

Our study sought to address this gap by elucidating how prolonged degeneration, specifically hAPP induced degeneration, affects olfactory glomerular circuitry at the ultrastructure level and how circuitry connectivity changes from baseline during degeneration and recovery. Our approach used serial electron microscopy to create the first known microcircuit reconstructions of the glomerular circuit, allowing us to analyze cellular composition, subcellular structures and connectivity complexity at baseline, after prolonged degeneration and during recovery.

## Materials & Methods

### Transgenic Lines

Transgenic mouse lines were obtained from Jackson Laboratories and maintained in a temperature and humidity-controlled room with 12-hour light/dark cycles in the Animal Facility at the National Institutes of Health. We used a tetO-hAPP line with the Swedish and Indiana familial AD mutations (Jankowsky, Melnikova, et al. 2005) (Jackson Laboratory #34846) and the OMP-TTA line (Jackson Laboratory #017754). OMP-hAPP animals were generated by crossing OMP-tTA with tetO-hAPP animals to make lines that selectively expressed hAPP in mature olfactory sensory neurons. Genotyping was done to identify animals that expressed both transgenes.

### PCR Primers

~~~
OMP-tTA: 5’ GGTTGCGTATTGGAAGATCAAGAGC 3’
         5’ GAGGAGCAGCTAGAAGAATGTCCC 3’
tetO-hAPP: 5’ CCGAGATCTCTGAAGTGAAGATGGATG 3’
         5’ CCAAGCCTAGACCACGAGAATGC 3’
~~~

### Turning Off hAPP Expression

Recovery animals were raised to 4 weeks of age, and then put on doxycycline chow (6g/kg) for 6 weeks, Doxycycline (DOX) prevents the binding of tTA to tetO, stopping hAPP expression. Animals in all conditions were perfused at 10 weeks.

### Electron Microscopy

Animals were transcardially perfused at 10 weeks old, with 10 mL PBS for one minute. Tissue was dissected and fixed in 20 mL fixative (2.5% glutaraldehyde, 4% PFA, 0.1 M Na cacodylate) for 48 hours. We selected areas for sectioning that were in the central glomerulus to avoid cell bodies. Tissue was then shipped to Renovo Neural for 3D-EM procedures.

### Volumetric Analysis

Volumetric analysis was conducted using Fiji, VAST Lite, MATLAB, 3D Studio Max. Each plane of the EM Z-stack was a separate image. These were then imported into VAST Lite and individual cellular processes were segmented. After segmentation each segmented cube was exported for reconstruction and volume measurements in MATLAB, using the VastTools package.

### Cell Identity Analysis

Olfactory sensory neurons (OSNs) were identified as dark processes, as previously described (Bourne and Schoppa 2017; Pinching and Powell 1971a). Light processes were assigned as either OBNs or glia according to their morphology and subcellular structures. Light processes with high numbers or densely clustered vesicles and/or the presence of postsynaptic densities were assigned as OBNs. Light processes that were not OBNs with spindly morphology and few vesicles were assigned as glia. Any remaining processes were classified as unassignable.

### Subcellular Analysis

All analysis on EM data was performed using Fiji and VastLite. Images were analyzed in 67-image stacks. Each image was cropped to 1014 pixels by 1014 pixels (5 microns by 5 microns). Postsynaptic densities, mitochondria, and apoptotic-like vacuoles are large structures that frequently spanned multiple images in the stack. Care was taken to avoid double-counting any structure. Sum totals of subcellular structures do not include the counts from unassigned processes.

### Postsynaptic Densities

Postsynaptic densities (PSDs) were identified and confirmed manually. PSDs were two adjacent neurons with a concentration of vesicles on the presynaptic neuron and a postsynaptic density on the postsynaptic neuron. Two categories of PSDs were considered: OSN to OBN synapses and OBN to OBN synapses.

### Mitochondria

Mitochondria were identified by the presence of an inner membrane with visible cristae. Mitochondria with an abnormal appearance (i.e. loose or disorganized internal cristae) were included in total counts, as this was an expected result of the introduction of hAPP (Ye et al. 2015).

### Apoptotic Vacuoles

Apoptotic-like vacuoles were dark structures with a circularity measurement (*circularity = 4pi(area/perimeter^2*)) above 0.50 and area equal to or greater than 0.020 microns squared. Vacuoles had to span at least two sections to be considered. Vacuoles were grouped as residing in an OSN, OBN or glia.

### Vesicles

Vesicles were counted by outlining intracellular space containing vesicles. The scorer only outlined vesicles with an area of less than .01 microns^2^ avoiding mitochondria, and hollow vesicles to create a region of interest (ROI). Multivesicular bodies were counted and given their own ROI. Vesicles were then counted by ImageJ using the ‘find maxima’ function with a noise tolerance of 48. Each ROI was saved with all maxima labeled and separately reviewed by another scorer. This scorer accounted for any missed regions, reviewed to make sure the appropriate structures were excluded and corrected any errors from the macro.

### Connectivity

Processes were tracked through the 67-image stacks and all synaptic connectivity was recorded manually (see PSD analysis). Processes involved in synapses were given a number according to their VAST Lite segmentation number. Connectivity diagrams were created using Microsoft PowerPoint to recreate the synaptic connectivity in each image stack. These diagrams were then used to compute number of PSDs per process, number of synaptic connections per process and path length.

### Statistical Analysis

The data on subcellular features are reported as means ± SEM. Normally distributed data was analyzed with the appropriate t-tests. Non-normal data was evaluated using the rank-sum method. Outliers were removed when appropriate using the ROUT method at Q=.1%. The *p* values are reported in figure legends and results (*\*p<.05 **p<.01 ***p<.001*). N indicates the number of processes analyzed.

## Results

We used OMP-hAPP transgenic mice to express hAPP specifically in mature OSNs, which send axons directly into olfactory glomeruli and are their only peripheral input. Each glomerulus is a complex neuropil containing multiple cell types centrally derived Olfactory Bulb Neurons (OBNs) such as Mitral/Tufted cells, Periglomerular cells and also glia (Figure 1A, B). To evaluate the ultrastructural effects of hAPP induced neurodegeneration on the glomerular circuitry we compared tissue samples from OMP-hAPP transgenic mice in three states: those expressing hAPP (Mutant); hAPP non-expressing transgenic mice (Control) and hAPP expressing mice followed by a recovery period (Recovery). Previous studies with the hAPP olfactory model showed that significant olfactory bulb and olfactory epithelium degeneration occurs by 3 weeks of age (Cheng et al. 2013; Cheng, Cai, and Belluscio 2011). Thus, to create a recovery condition, we put a subset of 4-week-old mutant mice on DOX chow, disrupting the binding of tTA and tetO, and halting the expression of hAPP (Cheng et al. 2013). Animals were kept on DOX until they reached 10 weeks of age, when tissue was taken for all three conditions (Figure 1C).

**Figure 1.**
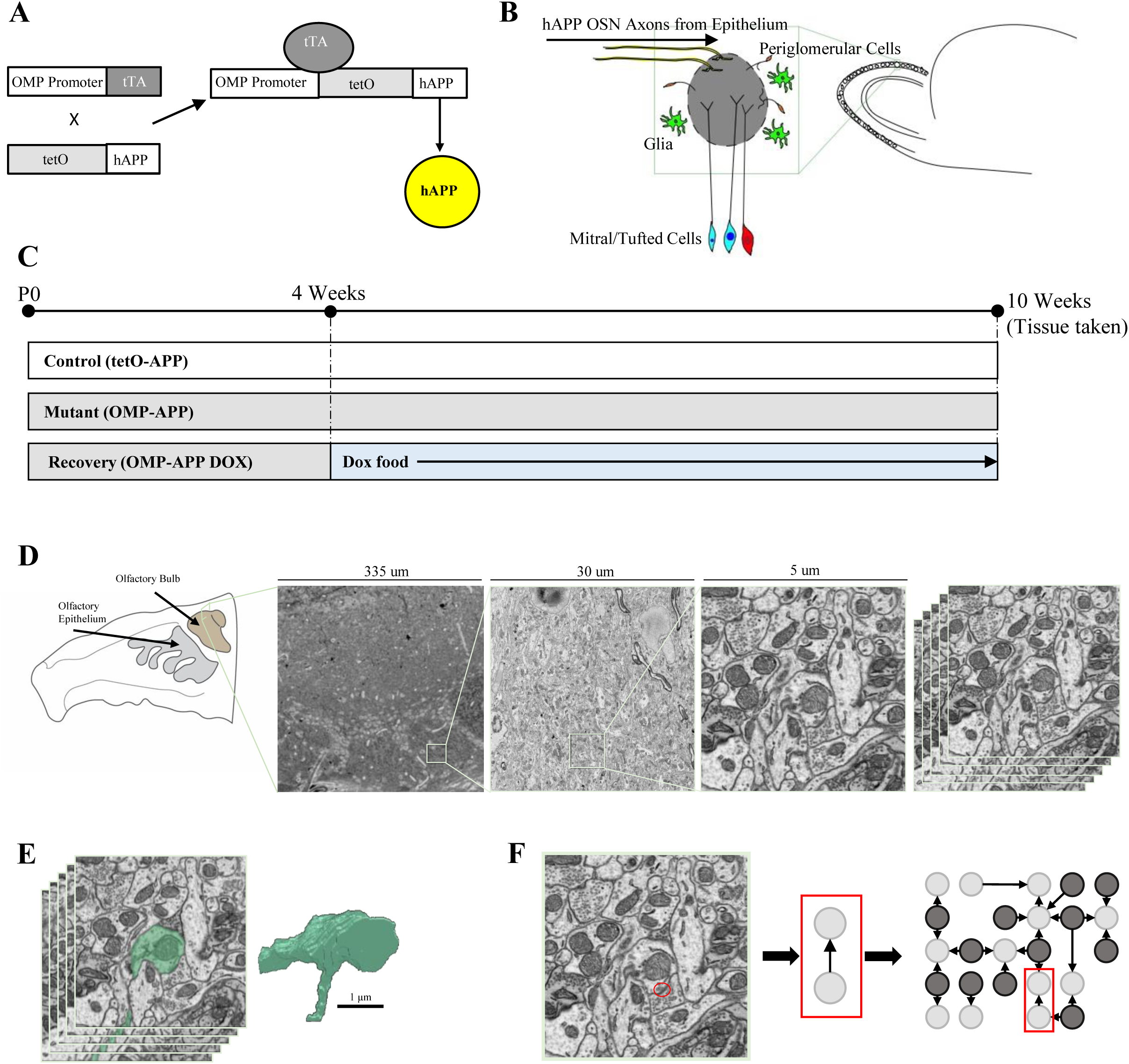
Study Design. **(A)** Diagram showing tetOff system used to generate hAPP induced degeneration and **(B)** visual representation of where hAPP would express in the olfactory system **(C)** Experimental time-lines. At 4 weeks the Recovery animal was put and kept on Dox food. All animals were sacrificed at 10 weeks. **(D)** Tissue was taken from the middle of the olfactory bulb, a low quality 335 um square image was used to select a 30 micron cube for high quality imaging from which a 5 micron cube was selected for consistency across conditions. **(E)** Cell processes were manually segmented and reconstructed using VastLite software. **(F)** Subcellular features were identified and quantified including post-synaptic densities (PSDs). Identified PSDs were assembled into connectomes.

Previous studies have shown degeneration and recovery of the olfactory bulb on the anatomical and behavioral level at this age. In this study we examined the subcellular effects of hAPP expression and recovery using serial section electron microscopy. We imaged a 35 microns^3^ region of central glomerular tissue from each condition then selected a 5-micron^3^ subregion for analysis of cellular composition and connectivity (Figure 1D, Figure Sup. 1). For each condition, cellular processes were first segmented and reconstructed allowing for volume measurements, process type identification, and quantification of subcellular structures (Figure 1E). One of the subcellular structures noted within each sample were PSDs, which were used to identify synaptic connectivity and recreate precise networks diagrams (Figure 1F).

### hAPP induction and blockage causes volumetric re-allotment of the cellular composition of the glomerulus

To measure the volumetric cellular composition of the glomerular tissue we used manual segmentation with a software interface to categorize cellular processes as either OSNs, OBNs, or glia (Figure 2A, B). Since previous studies have demonstrated that OSN projections appear darker in EM micrographs compared to surrounding cells (Au, Treloar, and Greer 2002; Pinching and Powell 1971b), it was relatively simple to identify OSN projections. We found that in the control condition OSNs contributed to 25.4% of the total volume in our sample, while they disappeared completely in the mutant condition, and reappeared at 41.5%, in the recovery animal (Figure 2C). Glial processes identified morphologically, were present in all three conditions and were most abundant in the mutant condition at 41.1%, compared to control animal at 25%, the increase may be a response to degeneration (Figure 2C). The volume of glial processes dropped substantially to 8.6% in the recovery condition (Figure 2C). The volume of OBN processes, identified through their lighter shading, were proportionally higher in the mutant at 49.5% while OBN volumes in the control and recovery mice were stable at 41.2% and 43.4% respectively (Figure 2C). A small, unassignable group was also present in all three conditions (Figure 2C).

**Figure 2.**
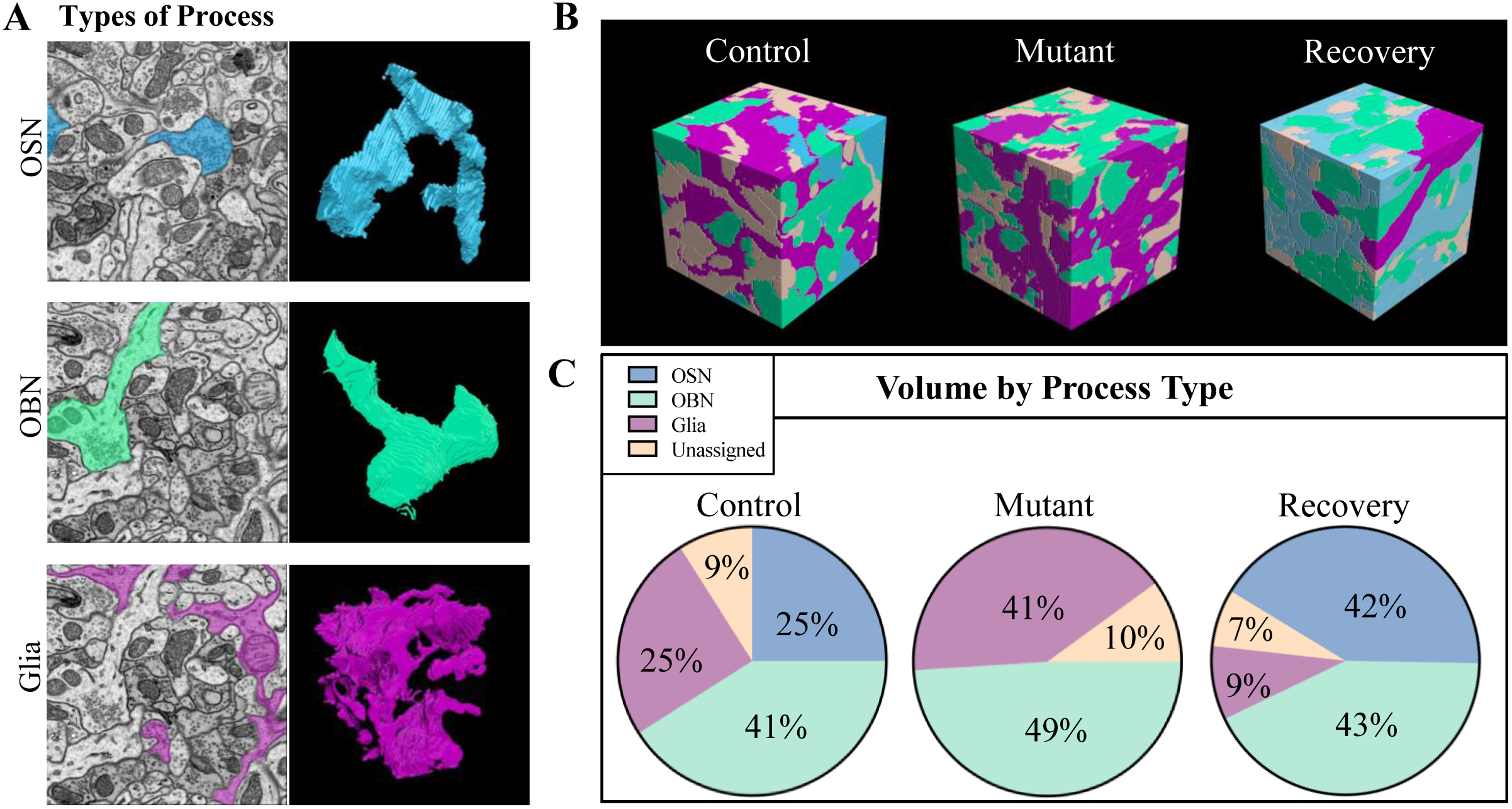
Volumetric reconstruction analysis. **(A)** Representative OSN, OBN and glia respectively showing the morphology of each process type. **(B)** Control, Mutant and Recovery connectome cubes, color coded by cell type. **(C)** Quantification of volumetric cellular composition in each condition. The volume of OBNs increase slightly in the Mutant condition (Control: 51.68 um^3^, Mutant: 62.00 um^3^, Recovery: 54.40 um^3^). The volume of glia increase dramatically in the Mutant condition and then decrease in the Recovery animal (Control: 31.31 um^3^ Mutant: 51.50 um^3^ Recovery:10.79 um^3^). OSNs disappear in the Mutant condition and reappear in the Recovery (Control: 31.89 um^3^ Mutant: 52.04 um^3^). The volume of unassignable processes remains relatively consistent across conditions (Control: 10.45 um^3^ Mutant: 11.82 um^3^ Recovery: 8.10 um^3^).

### hAPP induction causes subcellular changes in the olfactory glomerulus

To examine how subcellular structures responded to degeneration and recovery we identified and analyzed mitochondria and apoptotic like vacuoles across animal conditions (supplemental data). The numbers of mitochondria in the control (144) and mutant animals (132) were similar, while the recovery animal had the fewest mitochondria (109) (Figure 3A). To control for differences in cell volume, we measured the density of these features in each process type (Figure 3B). OSN mitochondrial density showed a reduction when comparing control and recovery conditions (*p=.0386*) and there were more processes without any mitochondria in the mutant, while glia mitochondrial density remained consistent across all conditions (control vs. mutant, *p =.2531*; control vs. recover, *p = .7167;* mutant vs recovery; *p= >0.999*). We also found no significant difference in OBN mitochondrial density across conditions although the number of processes with zero mitochondria was higher in the mutant as compared to control and recovery. When we compared the sum total of mitochondrial density in the entire tissue, across all three conditions we found no change between control and mutant or recovery and mutant (*p=.1830, p=.1967*, respectively), however, there was a significant decrease in density between control and recovery (*p=.0022*).

**Figure 3.**
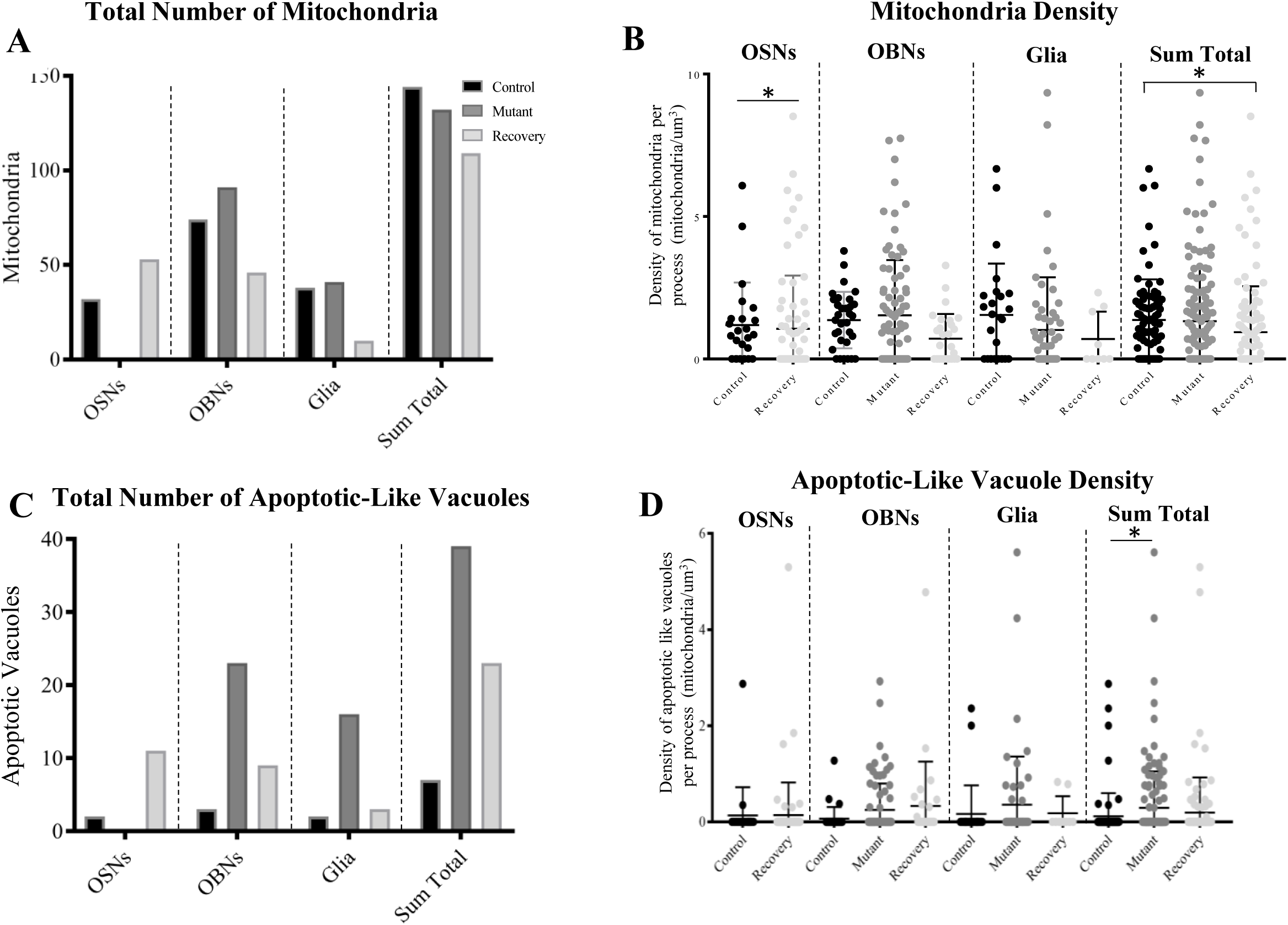
Identification and quantification of subcellular structures. **(A)** Number of mitochondria in each condition by process type. The Control animal had 144 mitochondria (OBN=74, OSN=32, Glia=38), the Mutant animal had 132 (OBN=91, Glia=41) and the Recovery animal had 109 (OBN=46, OSN=53, Glia=10). **(B)** Density of mitochondria in each condition by process type. OSN (Control vs Recovery, p = .0386; Control *n*=24, Recovery *n*=75). OBN (Control vs Mutant, p=.9117; Control vs Recovery, p=.0715; Mutant vs Recovery, p=.2713; Kruskal-Wallis statistic = 5.233; Control *n*=31, Mutant *n*=86, Recovery *n*=29.) Glia (Control vs Mutant, p=.2531; Control vs Recovery, p=.7167; Mutant vs Recovery, p=>.9999; Kruskal-Wallis statistic=2.031; Control *n*=26, Mutant *n*=58, Recovery *n*=9). Sum Total (Control vs Mutant, p=.1830; Control vs Recovery, p=.0022; Mutant vs Recovery, p=.1967; Kruskal-Wallis statistic=11.49; Control *n*=81, Mutant *n*=144, Recovery *n*=112). **(C)** Number of apoptotic like vacuoles in each condition by process type. The Control animal had 7 vacuoles (OSN=2, OBN=3, Glia=2), the Mutant animal had 40 (OBN=24, Glia=16) and the Recovery animal had 23 (OSN=11, OBN=9, Glia=3). (**D**) Density of apoptotic vacuoles in each condition by process type, there was no significant difference in cell type specific vacuole density, there was a significant increase in overall vacuole density Mutant processes when compared with Control *(p*=.0221).

Since changes in mitochondrial density might reflect an increase in cellular stress, we sought to investigate if there was evidence of degeneration at a subcellular level. To assess this, we identified and quantified subcellular apoptotic vacuoles. We found that indeed there were more vacuoles in the mutant than in the control and that the number of large vacuoles stayed elevated in our recovery model (7 vacuoles in control, 39 vacuoles in mutant, 23 vacuoles in recovery) (Figure 3C). Although we found no cell specific difference in vacuole density we did observe a significant increase in overall vacuole density between control and mutant tissue (*p=.0221*) (Figure 3D), indicating that hAPP expression may be producing a local effect to the glomerular circuitry.

To determine how hAPP induced degeneration and recovery affect synaptic machinery and subcellular transport we examined vesicle number. Interestingly, the total vesicle number within the entire tissue volume was similar across conditions, although it is difficult to precisely identify the vesicle types. Vesicles in our neuronal processes are most likely synaptic vesicles particularly when located opposite a post-synaptic density (PSD), while glial may contain vesicles with a range of neuroactive substances (Zorec et al 2016) and most non-neuronal vesicles are likely, transport vesicles. Our findings show a small increase in total vesical number in glia and found no significant change in vesicle density, indicating that degeneration does not impact the presence of vesicles in glia. We then turned to the presence of vesicles in our neuronal populations, which are largely synaptic vesicles (De Robertis and Franchi 1956; Doroquez et al. 2014). Here we found that the density of vesicles in mutant OBN processes was significantly higher than control or recovery OBNs (*p=.0012, p=.0030*, respectively) and there was a small, significant decrease in vesicle density in recovery OSNs (*p=.0132*) (Figure 4B).

**Figure 4.**
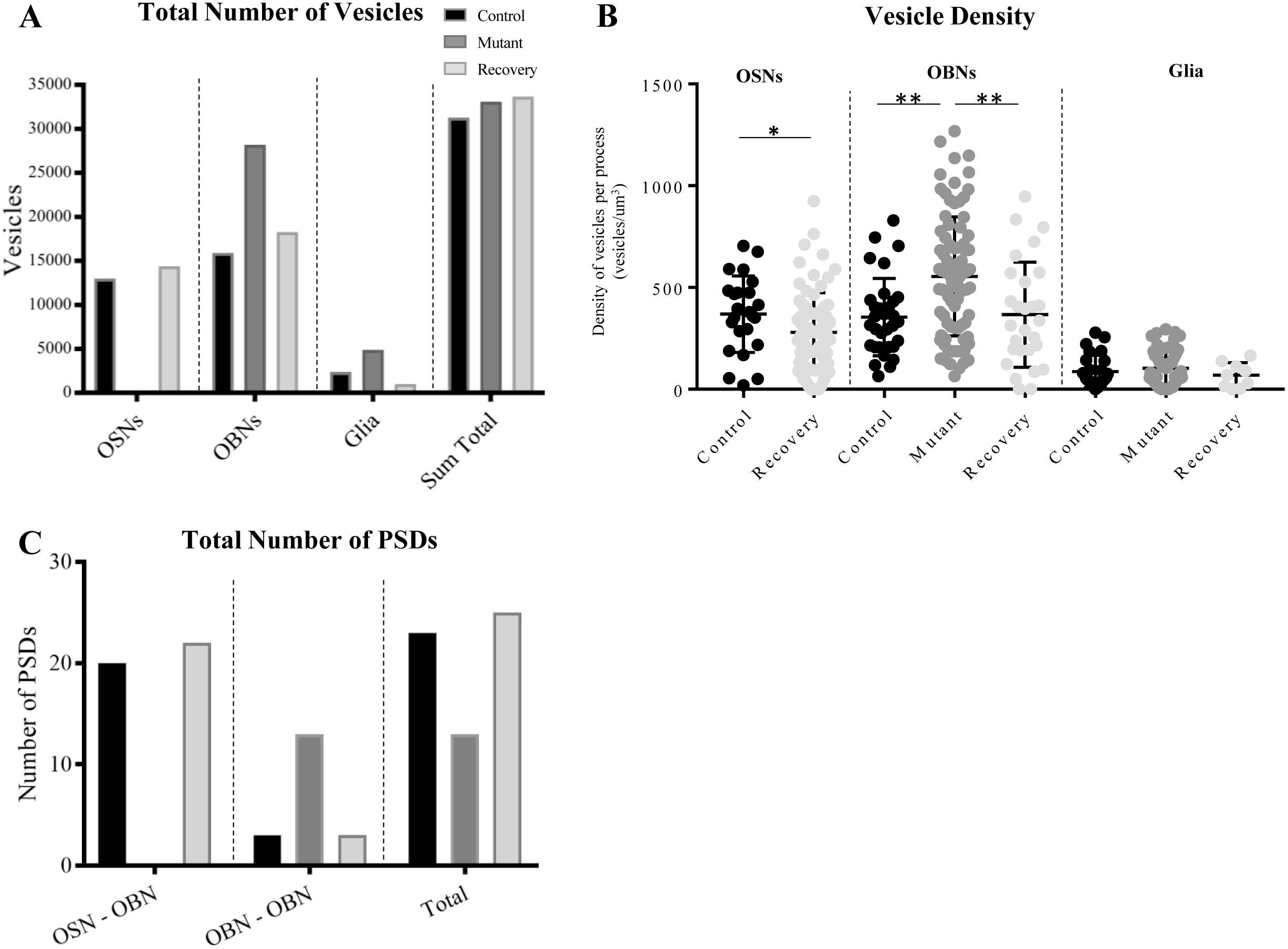
Subcellular synaptic machinery. **(A)** Number of vesicles in each condition by process type: OSNs (Control =12,984, Recovery=14,398), OBNs (Control= 15,914, Mutant=28,203, Recovery=18,265), Glia (Control=2,388, Mutant=4,897 Recovery=1,003), and sum total for each condition (Control =31,286, Mutant=33,100, Recovery=33,666). (**B**) There was a significant decrease in OSN vesicle density (p=.0132) or Glia density (p=.4623 F(2, 88) = .7783) however, *post-hoc* analysis revealed a significant change in vesicle density in OBNs between Control and Mutant (p=.0012) and Mutant and Recovery (p=.0030). (**C**) PSDs by type in each condition. OSN-OBN PSDs (Control=20, Recovery=22) OBN-OBN(Control=3, Mutant=13, Recovery=3).

We then asked if changes in neuronal vesicle density correlated with changes in connectivity. To analyze connectivity, we examined post-synaptic densities (PSDs). We found that the number of PSDs decreased in our mutant condition and rebounded to control levels in the recovery (Figure 4C). The number of OBN to OBN synapses increased in the mutant condition likely due to several changes; 1) the increase in total OBN volume in the mutant condition, and 2) the loss of OSNs creating more opportunity for OBN to OBN synapses.

### Synaptic connectivity can be partially recovered after hAPP induced loss

Using PSD data, we created connectomes for each tissue sample, allowing for further analysis of synaptic connectivity at each state (Figure 5A). We first determined the number of PSDs in each process that contained PSDs (Figure 5B). We found that mutant and recovery animals displayed a substantial reduction in the number of processes with more than one PSD as compared to control, however it is notable that the recovery condition had more processes with PSDs overall, indicating that neurons in the recovery condition may be in a growth state, actively seeking connections but with less complexity than controls. We next analyzed the total number of synaptic connections, both pre-and postsynaptic, for each process. Only 19% of processes in the mutant animal had more than one synaptic feature per process while 58% of processes in the control tissue and 52% of processes in the recovery tissue had more than one synaptic feature (Figure 5C). The recovery tissue does seem to robustly re-establish synaptic connectivity; however, it does not completely return to control levels. In the control animal 26% of processes had three or more synapses, while in the recovery condition 17% of processes had three or more synapses. We also examined path length in the connectomes and found all paths in the mutant condition were only one synapse long, while 20% of the paths in the control condition and 10% of paths in the recovery condition were two or three synapses long, indicating again a loss of complexity in the mutant and a partial regain of complexity in the recovery condition (Figure 5D). It is interesting to note that there were more overall processes in our recovery connectome than in either of the other conditions, perhaps indicating an overgrowth of newly generated OSN axons.

**Figure 5.**
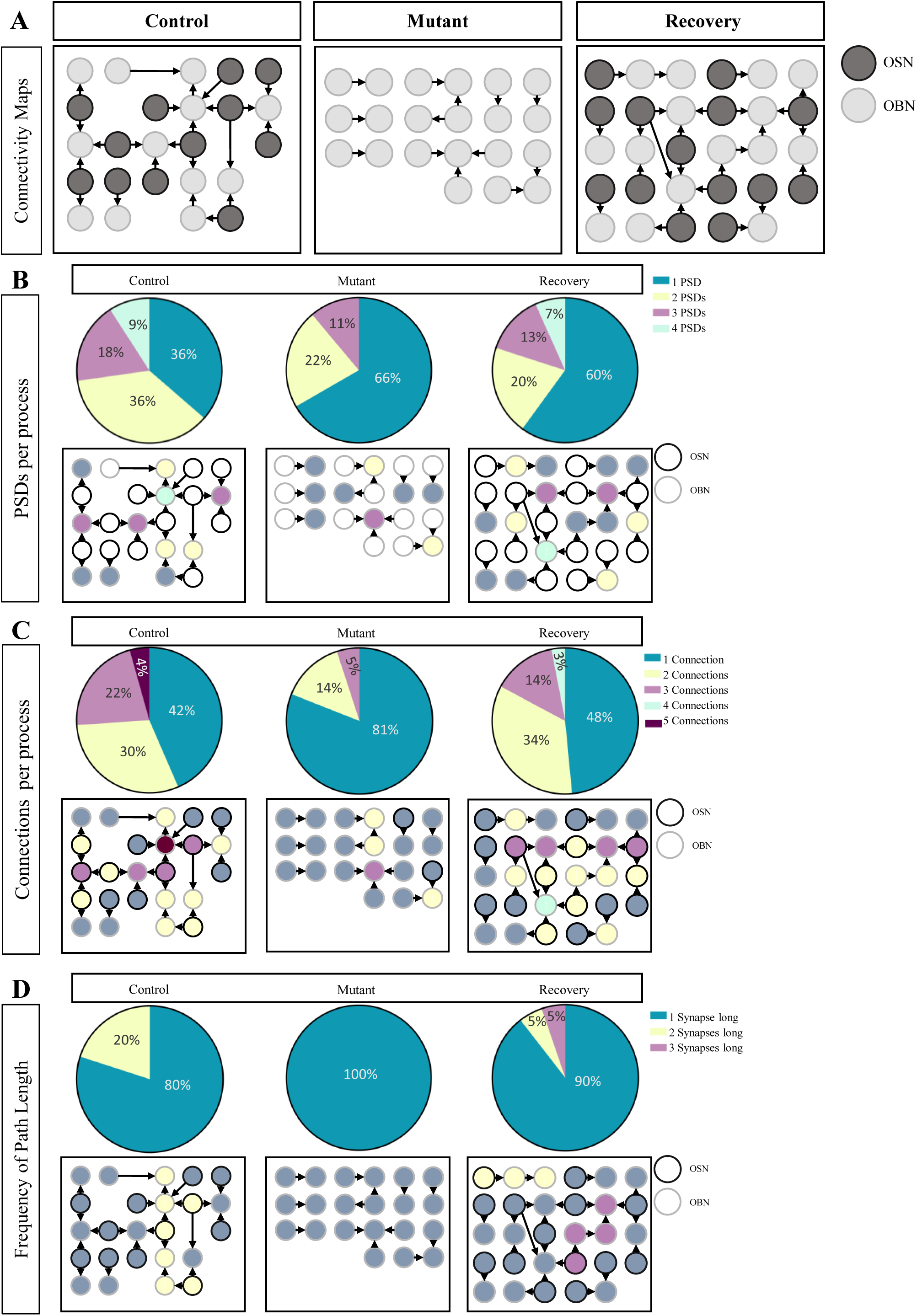
Glomerular Connectomes **(A)** Connectomes for each condition. Arrows indicate directionality of the synapse. Light cells are OBNs, dark cells represent OSNs. **(B)** Proportional representation of number of PSDs per process and corresponding connectome. Colors correspond with number of PSDs/cell **(C)** Proportional representation of how many synapses, either incoming or outgoing, per process and corresponding connectome. Colors correspond with number of synaptic connections/cell **(D)** Proportional representation of how frequently given path lengths occur and corresponding connectome. Colors correspond with the length of the path.

## Discussion

Ongoing OSN neurogenesis makes it possible to study circuit recovery following neural injury or degeneration (Cheng, Cai, and Belluscio 2011; Steuer, Schaefer, and Belluscio 2014). We found that hAPP-induced loss of OSNs broadly disrupts glomerular organization and yet partial recovery occurs when hAPP production is stopped suggesting that circuitry is not irreversible damaged. To evaluate changes in glomerular circuitry in three different states; before, during, and after hAPP-induced neurodegeneration, we performed serial EM reconstruction on representative portions of glomerular tissue in each state, taken from the center of the glomerulus and from similar regions of their OB. While we sought to be consistent in our tissue selection, it is important to note that the level of glomerular homogeneity has not been fully established, thus samples may contain selection bias. Nevertheless, this study represents the first 3D ultrastructural data showing how OSNs reestablish and rewire their axons in the glomerulus after disruption in a disease model. We find a clear ability for regenerating neurons to reconnect with downstream targets, demonstrating the strength of this system as a cellular model to study the effects of hAPP on circuitry and the subsequent ability to recover.

### hAPP induction causes reversible glomerular degeneration

We observed clear evidence of OSN degeneration through loss when hAPP was expressed. Our volumetric analysis also revealed a large increase in glial volume in the mutant condition, although it is unclear if this response is due to hAPP induced degeneration of the tissue or if glia, a highly mobile cell type, are simply occupying a higher proportion of the structure to compensate for the loss of OSNs. However, we suspect the large increase in glial volume we see in the mutant is due to degeneration caused by hAPP, as both microglia and astrocytes are known to respond to cell death as well as hAPP production (Doorn et al. 2014; Hovakimyan et al. 2013; Wirths et al. 2010). The significant increase in apoptotic vacuole density is further evidence of degeneration as the appearance of apoptotic vacuoles is considered a marker for neurodegeneration (Yang et al. 2005). Specifically, the rise in total number of vacuoles we see in OBNs in the mutant condition suggests that hAPP is having a non-cell autonomous effect. Interestingly in the recovery model the glial volume drops dramatically, and the density of apoptotic vacuoles decreases significantly suggesting the cellular degeneration in the mutant condition resolves when the cells are allowed to recover.

### Volumetric re-allotment of OSNs after hAPP induction

We observe a complete loss of OSNs in the mutant condition, however after recovery we noticed striking overgrowth of OSN axons reflected by increased proportion of OSN volume in recovery tissue. The loss of OSNs is consistent with previous IHC work on OMP-hAPP animals (Cheng et al. 2013; Cheng, Cai, and Belluscio 2011) although few studies have examined the reinnervation of regenerating OSN axons at the EM level. We reveal a volumetric imbalance within the glomerulus with OSN axons in the recovered condition occupying a larger proportion of the tissue as compared to controls indicating an exuberant regrowth. The overgrowth is reminiscent of initial glomeruli formation in perinatal animals, where OSN axons branch broadly and are refined as the animal ages (Marcucci et al. 2011; Potter et al. 2001). Because we drive hAPP expression from conception, our mutant animals do not establish the glomerular map at a normal developmental stage. Thus, our recovery animals may resemble this immature state due to the fact that they must establish and refine their glomerular maps beginning at 4 weeks of age when hAPP expression is turned off.

### hAPP induced cell loss alters the density of intracellular features

We asked if the degeneration and recovery we observed at the cellular level was reflected in the neuronal components of our model at the ultrastructural level by examining mitochondria and vesicles, features known to both respond to cellular degeneration and correlate with synaptic connectivity (Coleman and Yao 2003; Fond and Ravichandran 2016; Graziadei and DeHan 1973; Reddy et al. 2010; Sheng 2014; Tinari et al. 2008).

The overall increase in apoptotic vacuoles we observed in the mutant may be a consequence of the increase in OSN cell death within the glomerular volume. Furthermore the small reduction of apoptotic vacuole density e and high number of processes without mitochondria in the control and recovery condition suggests that mitochondrial density correlates with apoptosis and possibly cellular stress, an observation consistent with previous findings (Sheng 2014). Interestingly, we observed a large increase in vesicle density in the mutants, suggesting that whatever detrimental effects degeneration has on OBNs does not affect the ability of OBNs to make synaptic connections. Indeed, this increase in vesicle density pairs with an increase in OBN to OBN PSDs in the mutant condition suggesting that when OSN to OBN connectivity is eliminated in the mutant conditions, there may be a compensatory increase in OBN to OBN activity. Interestingly OSNs appear more stable with both mitochondrial density, vesicle density and overall PSD number. However, we did observe a small but significant decrease in vesicle density in the recovery condition suggesting that OSN axons while present in the recovery condition may not be synaptically mature.

### Degeneration-induced loss and subsequent recovery of connectivity

The theory of OSN overgrowth in the recover condition was supported by our analysis of connectomes. In the recovery model the number of synaptic connections per cell reached near control levels and longer path lengths reappeared. The number of PSDs per process did not fully return, however we saw a small increase in overall PSD number compared to control, supporting the idea that OSN axons in the recovery state are innervating glomeruli and actively forming new connections. During degeneration we found decreased connectivity including reductions of PSDs, synaptic connections per process, as well as path lengths, consistent with previous observations in hippocampal neurons coculture experiments that mutant hAPP decreases synaptic connectivity (Katsurabayashi et al. 2016). Previous studies report that following OSN lesioning, regenerating OSN axons can retarget the OB, albeit in a disorganized manner but, these reports did not address whether new OSNs can re-establish their connection and how it effects the OB circuits (St John and Key, 2003; Costanzo, 2000). Our data suggest that despite the anatomical disorganization associated with a disrupted glomerular map, the regenerating OSN axons can indeed establish complex synaptic connections after disruption. Moreover, they are essential for longer path lengths within the glomerular circuitry suggesting that they play a bridging role to enable broader network communication.

Together, these data suggest that the rodent olfactory glomerulus can largely recover connectivity after OSN cell death and subsequent regeneration. The sample bias in this initial study is important to consider yet our analysis has enabled some predictive insights into glomerular network connectivity and proportional distribution of cell types and subcellular structures that future EM reconstructive efforts can build upon. In addition, studies of glomerular homogeneity both in adults and during development will be important for understanding the generalizability of any reconstructive findings. As the cellular mechanism of APP associated degeneration is still was not well understood this study adds a new perspective and platform for understanding network disruption subsequent restoration as well as the underlying mechanism.

## Acknowledgements

We would like to thank Waverley He and Aakash Basu for assistance with software implementation and segmentation protocols. We thank Renovo Neural Inc. (Cleveland, Ohio) for assistance with performing the serial-EM image acquisition.

**Figure Supplemental 1.**

EM Stacks. 5 micron^3^ stack for each condition.

**Figure Supplemental 2**

Counts of subcellular features. Spread sheet corresponding to each image of a stack for each condition with the final sheet being a tally of all images for each condition.

